# Disinhibitory and neuromodulatory regulation of hippocampal synaptic plasticity

**DOI:** 10.1101/2020.10.13.337188

**Authors:** Inês Guerreiro, Zhenglin Gu, Jerrel L. Yakel, Boris S. Gutkin

## Abstract

Hippocampal synaptic plasticity, particularly in the Schaffer collateral (SC) to CA1 pyramidal excitatory transmission, is considered as the cellular mechanism underlying learning. The CA1 pyramidal neurons are embedded in an intricate local circuitry that contains a variety of interneurons. The roles these interneurons play in the regulation of the excitatory synaptic plasticity remains largely understudied. Our recent experiments showed that repeated cholinergic activation of α7 nACh receptors expressed in oriens-lacunosum-moleculare (OLMα2) interneurons could induce LTP in SC-CA1 synapses, likely through disinhibition by inhibiting stratum radiatum (s.r.) interneurons that provide feedforward inhibition onto CA1 pyramidal neurons, revealing a potential mechanism for local interneurons to regulate SC-CA1 synaptic plasticity. Here, we pair *in vitro* studies with biophysically-based modeling to uncover the mechanisms through which cholinergic-activated GABAergic interneurons can disinhibit CA1 pyramidal cells, and how repeated disinhibition modulates hippocampal plasticity at the excitatory synapses. We found that α7 nAChR activation increases OLM activity. OLM neurons, in turn inhibit the fast-spiking interneurons that provide feedforward inhibition onto CA1 pyramidal neurons. This disinhibition, paired with tightly timed SC stimulation, can induce potentiation at the excitatory synapses of CA1 pyramidal neurons. Our work further describes the pairing of disinhibition with SC stimulation as a general mechanism for the induction of hippocampal synaptic plasticity.

Disinhibition of the excitatory synapses, paired with SC stimulation, leads to increased NMDAR activation and intracellular calcium concentration sufficient to upregulate AMPAR permeability and potentiate the synapse. Repeated paired disinhibition of the excitatory synapse leads to larger and longer lasting increases of the AMPAR permeability. Our study thus provides a novel mechanism for inhibitory interneurons to directly modify glutamatergic synaptic plasticity. In particular, we show how cholinergic action on OLM interneurons can down-regulate the GABAergic signaling onto CA1 pyramidal cells, and how this shapes local plasticity rules. We identify paired disinhibition with SC stimulation as a general mechanism for the induction of hippocampal synaptic plasticity.

## Introduction

Since the discovery of long-term potentiation (LTP) in the hippocampus in 1973, this structure has been used as a model to study the mechanisms of synaptic plasticity induction (Bliss, 1973). The hippocampal SC to CA1 synapses are among the most studied for synaptic plasticity. The majority of the work has focused on excitatory glutamatergic transmission. However, recent work has shown the importance of inhibitory inputs in the modulation of local hippocampal synaptic plasticity (Saudargiene & Graham, 2015; Chevaleyre & Piskorowski, 2014; Wigstrom & Gustafsson, 1985).

The hippocampal network comprises a large variety of locally connected GABAergic interneurons exerting powerful control on network excitability. GABAergic interneurons receive important cholinergic innervation from the medial septum and are endowed with a variety of different subtypes of nicotinic acetylcholine receptors (nAChRs) that are thought to be involved in regulating excitability, plasticity, and cognitive functions (Griguoli & Cherubini, 2012; Levin, 2002; Yakel, 2012). However, the mechanisms through which cholinergic-activated GABAergic interneurons modulate excitatory transmission and induction of hippocampal plasticity at excitatory synapses remain largely unclear.

Alterations of cholinergic action on hippocampal GABAergic interneurons have been implicated in cognitive dysfunction in Alzheimer’s disease (AD) (Schmid, et al., 2016). Understanding how cholinergic activation of GABAergic interneurons regulates hippocampal excitability and induces synaptic plasticity will provide insight for understanding higher brain functions, namely memory impairment, with potential relevance for AD treatments.

Previous studies showed that activation of OLMα2 interneurons increases SC to CA1 transmission, likely through disinhibition by reducing the activity of stratum radiatum (s.r.) interneurons that in turn provide feed-forward inhibition onto pyramidal neurons (Leão, et al., 2012). Consistent with these studies, Gu et al. found that activation of OLMα2 interneurons increased SC to CA1 EPSCs and reduced IPSCs (Gu, et al., 2020). However, the mechanisms through which the activation of the inhibitory interneurons OLMα2 is regulating the activity of inhibitory interneurons targeting the CA1 pyramidal cell, and how this facilitates the potentiation of SC-evoked EPSPs of the CA1 pyramidal cells, remain elusive.

In this work, we use a minimal biophysical circuit model, driven quantitatively by in vitro data, to examine how activation of OLM cells modulates the activity of fast-spiking interneurons whose GABAergic inputs are co-localized with the SC glutamatergic synapses onto a CA1 pyramidal cell dendrite, and how this promotes the induction of plasticity at the SC-CA1 synapse. More specifically, we seek to determine how cholinergic activation of the OLM cells through the post-synaptic α7 nAChRs can down-regulate the GABAergic signaling onto the pyramidal cells, and how a decrease in inhibitory inputs to the pyramidal cells can modulate the plasticity of the excitatory SC-CA1 synapse. We thus constructed a minimal circuit consisting of a single compartment spiking model of an OLM interneuron with postsynaptic α7 nAChRs, a fast-spiking interneuron with postsynaptic AMPA and GABA_A_ receptors, and a pyramidal cell dendritic compartment with postsynaptic AMPA, NMDA and GABA_A_ receptors, connected as schematically shown in Figure 1.

**Figure 1:**
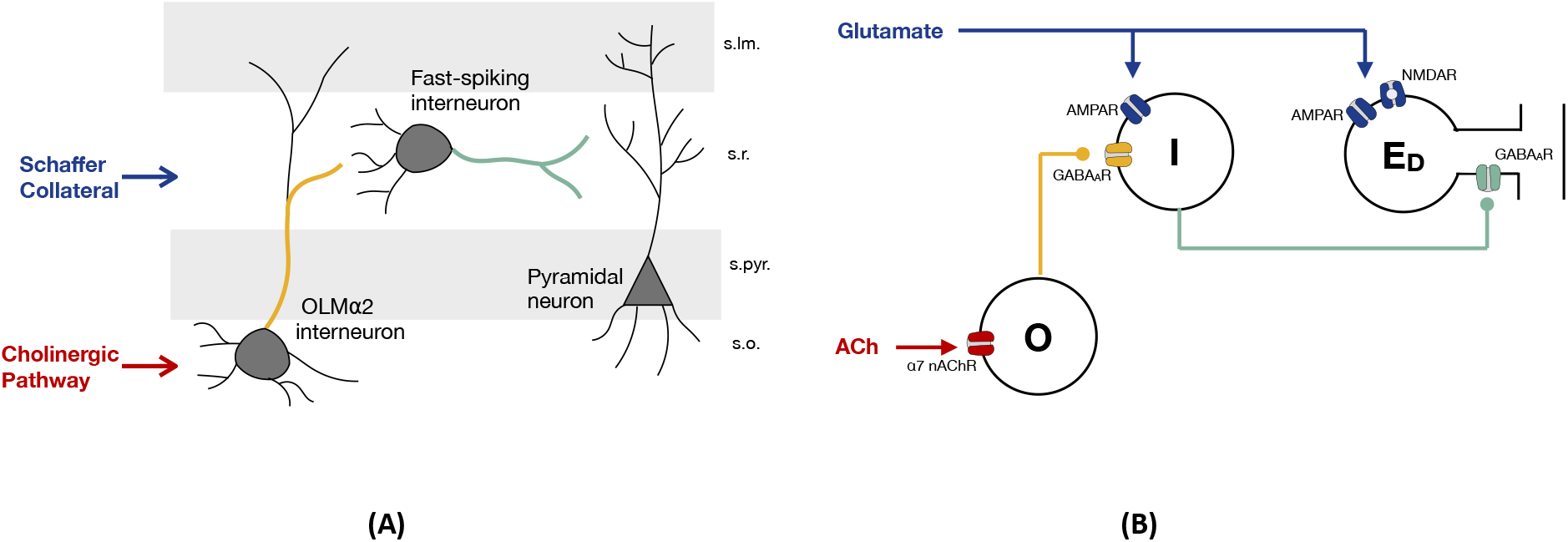
Disynaptic disinhibition circuit for ACh induced long-term plasticity in the CA1. **(A)** Simplified wiring diagram of an interneuron network that mediates feedforward inhibition in the CA1 region of the hippocampus. Activation of the Schaffer Collateral (SC) pathway leads to the activation of CA1 pyramidal cell dendrites and of the stratum radiatum (s.r.) interneurons, which provide feedforward inhibition onto the pyramidal cell. Cholinergic activation of OLMa2 interneurons in stratum oriens (s.o.) leads to the inhibition of the s.r. interneurons, counteracting SC feedforward inhibition (Leão, et al., 2012). **(B)** Minimal network to investigate plasticity induced by pairing of cholinergic and SC activation. Glutamate activates postsynaptic AMPA and NMDARs at the pyramidal cell dendritic compartment E_D_ and postsynaptic AMPARs at I-cells, which in turn provide feedforward inhibition onto E_D_ by activating postsynaptic GABA_A_Rs. Cholinergic inputs act on postsynaptic a7 nAChRs of O-cells, which results in GABA release of the O-cells that it is going to bind to postsynaptic GABA_A_Rs of the I-cells.

It is currently well accepted that LTP triggering processes require calcium influx through NMDARs. At the same time, LTP has been associated with changes in the properties of postsynaptic AMPARs; namely changes in their number and phosphorylation state (Barria, Muller, Derkach, Griffith, & Soderling, 1997; Collingridge, Kehl, & McLennan, 1983; Lüscher & Malenka, 1994). To reflect these mechanisms, we employ the calcium-based synaptic plasticity model (proposed by Shouval et al. (Shouval, Bear, & Cooper, 2002)) to model synaptic plasticity of the SC-CA1 excitatory synapse. We describe how a decreased release of the inhibitory neurotransmitter GABA could lead to increased NMDAR activation and intracellular calcium concentration, leading to changes in the CA1 pyramidal cell postsynaptic AMPA-mediated conductance. Our work also shows how nicotinic cholinergic action on OLM interneurons can downregulate the GABAergic signaling onto CA1 pyramidal cells and facilitate potentiation of the SC-CA1 synapse.

## Methods

### Animals and materials

All procedures related to the use of mice followed protocols approved by the Institutional Animal Care and Use Committees of the NIEHS. ChAT-cre mice (B6;129S6-Chattm2(cre)Lowl/J), Sst-cre mice (Ssttm2.1(cre)Zjh), and floxed *a7* nAChR knockout mice (B6(Cg)-Chrna7tm1.1Ehs/YakelJ) were originally purchased from Jackson Laboratory and then bred at NIEHS. OLMα2-cre mice (Tg(Chrna2cre)OE29Gsat/Mmucd) were originally obtained from Mutant Mouse Resource and Research Centers (MMRRC) and then bred at NIEHS. Mice (of either sex) were used for slice culture from day 6 to 8.

Culture media were from Sigma and Invitrogen. AAV serotype 9 helper plasmid was obtained from James Wilson at the University of Pennsylvania. The AAV vector containing floxed ChR2 (Addgene #20297) and floxed eNpHR (Addgene #26966) were obtained from Karl Deisseroth (Witten et al., 2010; Gradinaru et al., 2010). AAV viruses were packaged with serotype 9 helper at the Viral Vector Core facility at NIEHS.

### Brain slice culture and AAV virus infection

To study the effects of cholinergic coactivation on the plasticity of SC to CA1 synapses in Figure 2, coronal septal slices (350 fflm) from ChAT-cre mice and horizontal hippocampal slices from floxed *a7* nAChR mice or OLMα2-cre/floxed *a7* nAChR mice (350 fflm) were cut with Leica VT1000S vibratome. Medial septal tissue containing cholinergic neurons was then dissected out and placed next to the hippocampus on a 6-well polyester Transwell insert (Corning) and cultured there for about 2 weeks before being used for experiments, similar as previously described (Gu and Yakel, 2017). AAV viruses containing double floxed ChR2 construct (5 nl) were microinjected to the septal tissue with a micro injector (Drummond Scientific) on the second day of culturing. To study the effects of disinhibition on the plasticity of SC to CA1 synapses in Figure 4, horizontal hippocampal slices from Sst-cre mice were cultured and AAV viruses containing double floxed eNpHR construct were microinjected to the hippocampus the next day.

**Figure 2:**
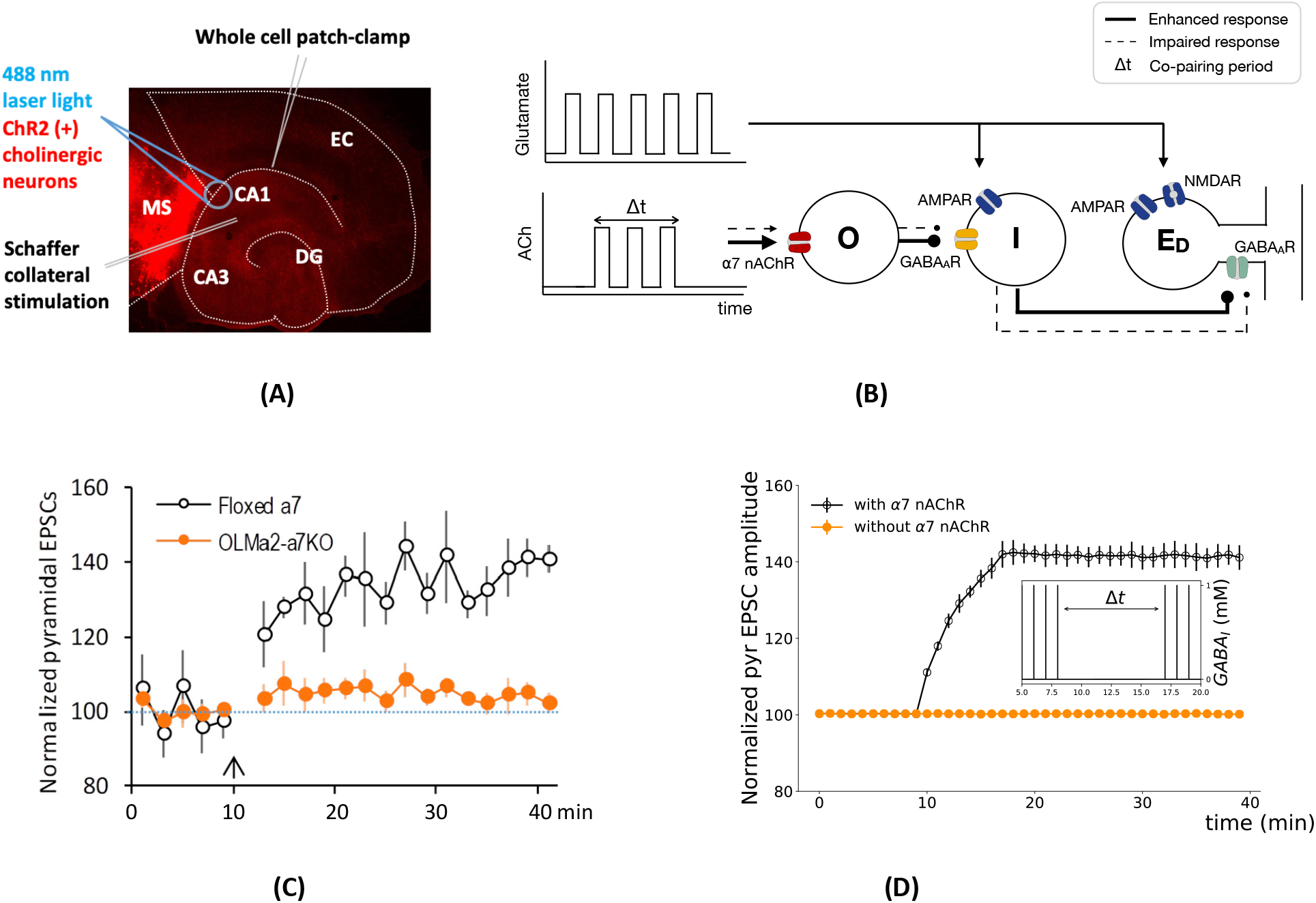
Cholinergic activation of OLM interneurons potentiates SC-evoked EPSCs. **(A)** Scheme of *in vitro* induction of cholinergic pairing induced hippocampal synaptic plasticity. EPSCs were recorded from CA1 pyramidal neurons. Cholinergic neurons were activated via channelrhodopsin-2 that was specifically expressed in ChAT-positive neurons. The Schaffer collateral (SC) pathway was activated by a stimulating electrode. Adapted from (Gu, Alexander, Dudek, & Yakel, 2017). **(B)** Scheme of minimal network used to study role of cholinergic inputs in the potentiation of SC-evoked EPSCs. Glutamatergic inputs activate the pyramidal cell dendritic compartment (E_D_) and the fast-spiking interneuron (I) that projects to it. The OLM interneuron (O) is activated by ACh during the co-pairing period. **(C)** Normalized SC-evoked EPSC responses from CA1 pyramidal neurons showing that the enhancement of EPSCs was impaired in hippocampal slices from mice with selective α7 nAChR knockout in OLMα2 interneurons. Adapted from (Gu, Alexander, Dudek, & Yakel, 2017). **(D)** Numerical simulation of normalized EPSC amplitude when glutamatergic inputs acting on the I-cell and E_D_ are paired with cholinergic inputs acting on the O-cell. The EPSCs are calculated as the sum of postsynaptic AMPA and NMDA currents, I_AMP A_ and I_NMDA_, and are a result of 10 simulations with white noise on the E_D_ membrane potential. Normalization of the results was calculated according with the expression (100 + (EPSC - EPSCmin).(150-100))/(EPSCmax - EPSCmin). Inset: Concentration of GABA released from fastspiking interneurons (I), calculated according to equation 15 (see Methods).

**Figure 3:**
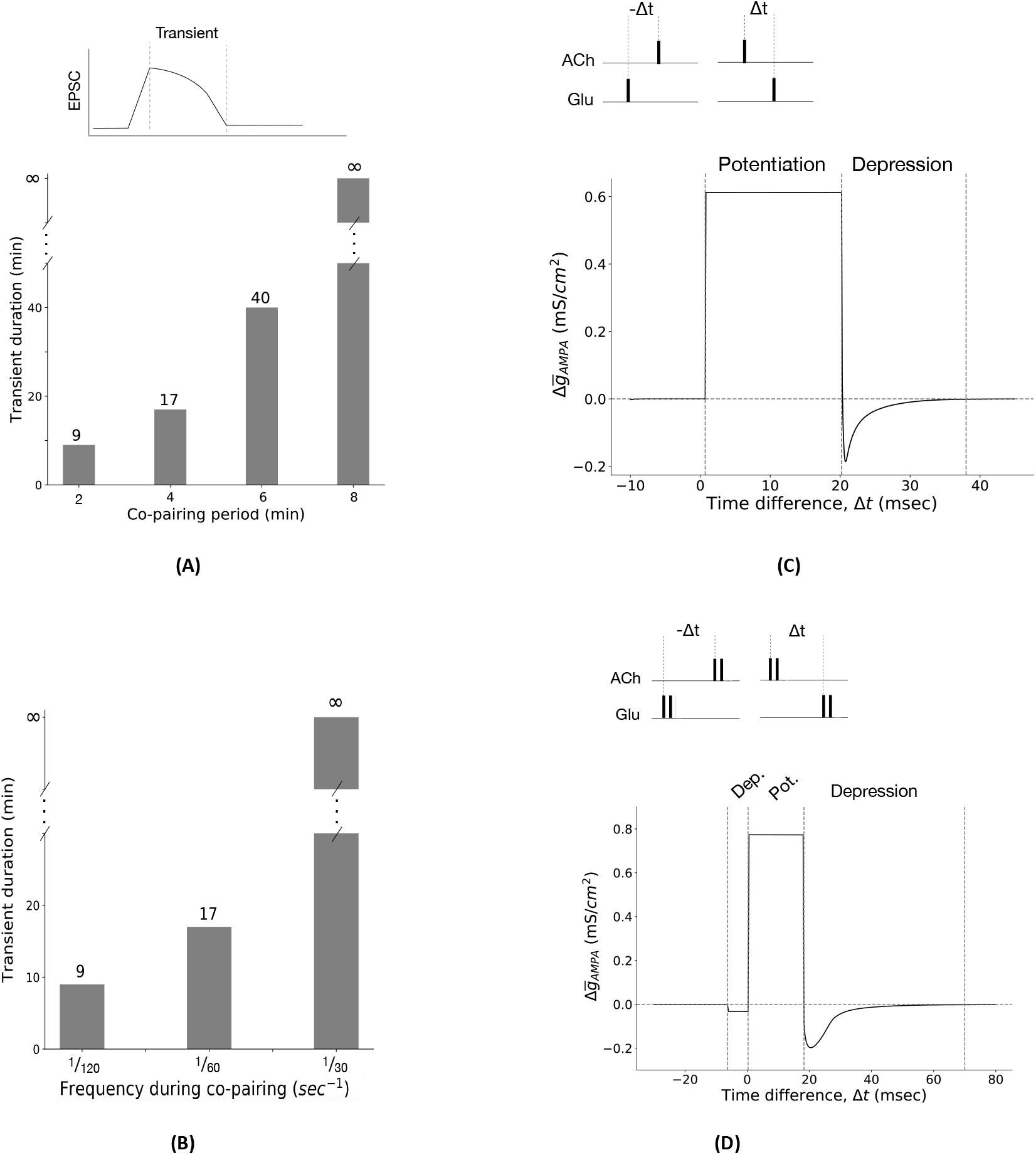
Co-pairing temporal parameters determine the duration and polarity of synaptic plasticity: relative timing between cholinergic and glutamatergic stimulation, extent of the co-pairing period, and the frequency of stimulation. **(A)** Synaptic strength transient duration is proportional to the extent of the pairing period. Here, the transient duration is defined as the time it takes the EPSC to go back to baseline after co-pairing is over. The I-cell and E_D_ receive a pulse of glutamate per minute. During the co-pairing period, the O-cell receives a pulse of ACh per minute, 1 msec prior to the pulses of glutamate. **(B)** Synaptic strength transient duration is proportional to the frequency of the ACh and glutamate pulses during the co-pairing period. Before and after co-pairing period, the I-cell and E_D_ receive a pulse of glutamate per minute. During the co-pairing period (4 minutes), the frequency changes to 1/20, 1/60 or 1/30 sec^-1^, and the O-cell receives a pulse of ACh 1 msec prior to the pulses of glutamate with the same frequency. **(C)** Relative paring timing provides a window of opportunity for plasticity. If glutamatergic inputs are administrated within 0.7*msec* < Δ*t* < 20.2*msec* the E_D_ excitatory synapse is potentiated. If glutamatergic inputs are administrated within 20.2*msec* < Δ*t* < 38*msec*, depression is induced. Negative values of Δ*t* correspond to glutamatergic stimulation prior to cholinergic activation. In that case, there are no changes in synaptic strength 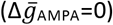. The change in the AMPAR conductance 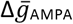 is measured 60 msec after one pairing. **(D)** Pairing multiple pulses of glutamate and ACh can change window of opportunity for plasticity. Two pulses of glutamate and ACh with a frequency of 1/5 msec^-1^ are paired. If glutamatergic inputs arrive within −6.28*msec* < Δ*t* < 0.3*msec* or 18.27*msec* < Δ*t* < 70*msec* of the cholinergic inputs, depression is induced. If glutamatergic inputs are administrated within 0.3 < Δ*t* < 18.27 *msec*, the E_D_ excitatory synapse is potentiated. The change in the AMPAR conductance 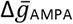 is measured 60 msec after one pairing. Parameters for subfigure (D): g_N_=0.18 mS/cm^2^ and g_G_=0.95 mS/cm^2^ for the postsynaptic receptors of E_d_. All other parameters are as indicated in the Methods.

**Figure 4:**
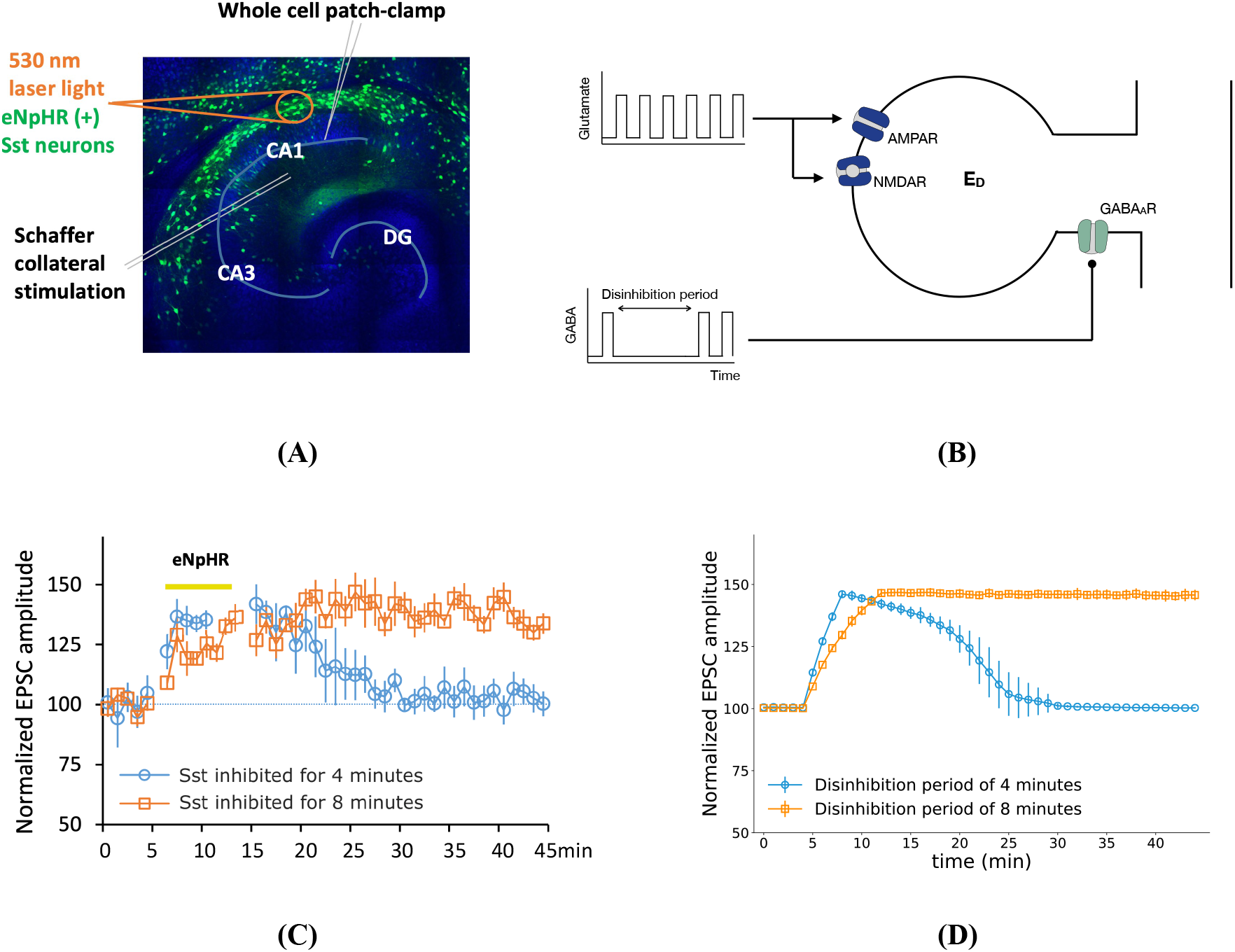
Disinhibition of CA1 pyramidal cell facilitates induction of hippocampal synaptic plasticity. **(A)** Scheme of in vitro induction of hippocampal synaptic plasticity through concurrent Sst inhibition. EPSCs were recorded from CA1 pyramidal neurons. Sst neurons were inhibited via eNpHR that was specifically expressed in Sst-positive neurons. The Schaffer collateral (SC) pathway was activated by a stimulating electrode. **(B)** Schematic representation of a CA1 pyramidal neuron dendritic compartment *E_D_* with post-synaptic GABA_A_, AMPA and NMDA receptors used to study the disinhibitory mechanisms for induction of plasticity at the SC-CA1 excitatory synapse. The dendritic compartment of the pyramidal cell receives one pulse of both Glutamate and GABA per minute, except during the disinhibition period where it only receives pulses of Glutamate. Glutamate binds to the excitatory AMPA and NMDA receptors, while GABA binds to the inhibitory GABA_A_ receptor. **(C)** Experimental measurements showing the effects of inhibition of Sst and OLM*α*2 interneurons in s.o. on SC-evoked EPSCs. Inhibition of Sst interneurons from t=4min to t=8min enhanced the SC-evoked EPSC amplitude of CA1 Pyramidal cell, followed by a return to the baseline after inhibition period (blue line). Inhibition of Sst interneurons from t=4min to t=12min resulted in the increase of SC-evoked EPSCs amplitude, which remained potentiated after inhibition period (orange line). **(D)** Numerical simulation of normalized EPSCs of *E_D_* for a disinhibition period of 4 minutes (from t=5 min to t=9 min) and 8 minutes (from t=5 min to t=13 min).

### Whole-cell patch clamp recordings

SC to CA1 excitatory post-synaptic currents (EPSCs) were recorded from hippocampal CA1pyramidal neurons under whole-cell patch clamp, similar as described before (Gu and Yakel, 2011, 2017). Briefly, 2-3 weeks after culturing, the slices were removed from transwell inserts and put into a submerged chamber, continuously perfused with 95%O2/5%CO2 balanced ACSF (in mM, 122 NaCl, 2.5 KCl, 2 MgCl_2_, 2 CaCl_2_, 1.2 NaH_2_PO_4_, 25 NaHCO_3_, 25 glucose) at room temperature. EPSCs were recorded at −60 mV under voltage clamp through a glass pipette filled with internal solution (in mM, 130 potassium gluconate, 2 MgCl2, 3 MgATP, 0.3 Na2GTP, 10 KCl, 10 HEPES, and 1 EGTA, with pH ~7.2-7.3 and osmolarity ~280-290 mOsm). Whole-cell patch clamp recordings were performed with Multiclamp 700B amplifier (Axon Instruments). Data were digitized with Digidata 1550, collected with Clampex. The amplitudes of EPSCs were analyzed with Clampfit and graphs were drawn with Excel. The amplitudes were normalized to the mean of the 10-min baseline recording before cholinergic pairing or disinhibition pairing. Values were presented as mean ± SEM.

EPSCs were evoked every 60 seconds by stimulating the SC pathway with an electrode placed in the stratum radiatum through a stimulator (Grass S88X). The stimulation intensity was 1-10 μA for 0.1 ms. To study the effects of cholinergic coactivation on SC to CA1 synaptic plasticity in Figure 2, cholinergic terminals in the hippocampus were optogenetically activated (10 pulses at 10 Hz, 1 sec before SC stimulation) through ChR2 that was selectively expressed in ChAT-cre positive (cholinergic) neurons. ChR2 was activated with 488-nm laser light (20 ms) through a 40× objective with an Andor spinning disk confocal microscope (Andor technology). To study the effects of disinhibition on SC to CA1 synaptic plasticity in Figure 4, Sst positive neurons were inhibited optogenetically through eNpHR which was activated with 530-nm laser light for 1 sec over SC stimulation.

### Model

The minimal network used in this study consists of an OLM cell (O), a fast-spiking interneuron (I), and a pyramidal cell. All the cells in the network are modeled as point neurons. Since we are interested in studying local changes at the SC-CA1 synapse, the pyramidal cell is represented by a dendritic compartment (E_D_).

### Neuron dynamics models

The O and I cells are modeled using the standard Hodgkin-Huxley currents (Hodgkin & Huxley, 1952) (transient INa, delayed rectifier potassium I_K_, and leak I_leak_), with synaptic currents I_Syn_. Its membrane potential V_m_ is described as follows:

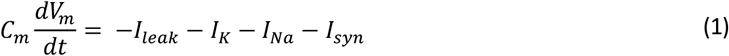

where Cm is the membrane capacitance. The I_leak_, I_k_ and I_Na_ currents are given by:

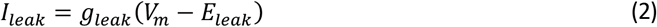

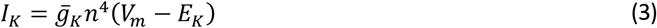

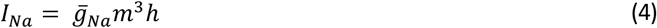

here 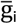 and E_i_ are, respectively, the maximal conductance and reversal potential of channel i (i=leak, K, Na), and m, h and n are gating variables that obey the following differential equation:

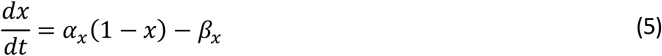

where α_x_ and β_x_ are voltage-dependent rate constants.

In addition, the O-cells have an applied current I_app_ = −2.6 μA/cm^2^, a persistent Na-current I_p_, and a hyperpolarization-activated inward current I_h_ (with a slow and fast component), as described in (Rotstein, et al., 2005):

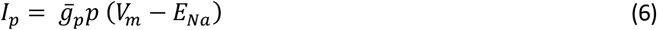

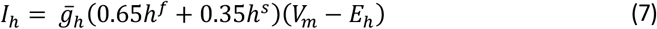

While the gate variable p obeys equation (5), h^f^ and h^s^ are described by the following equation:

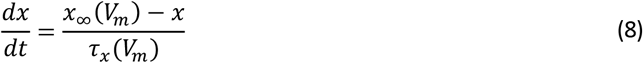

where x_∞_ is the voltage dependent steady state and τ_x_ the time constant. Definitions for the *α_x_, β_x_, x*_∞_ and *τ*_x_ for each of the dynamic variables are as follows.

For the O-cells:

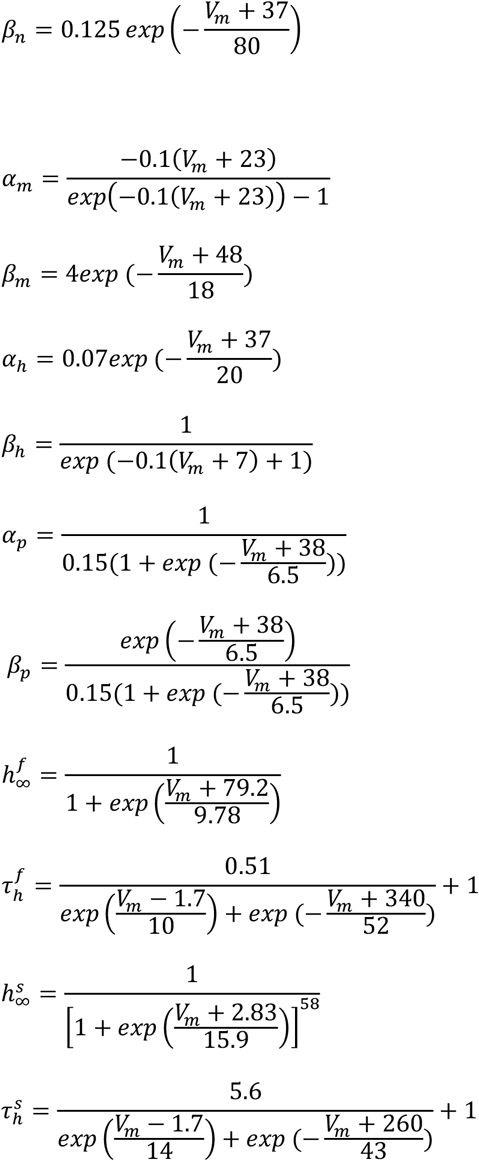

For the I-cells:

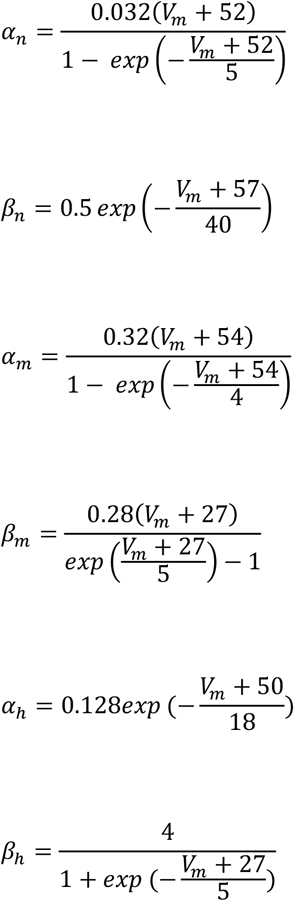

The parameter values used in the simulations are the ones presented in Table 1.

**Table 1:** Parameters of pyramidal cell, OLM interneuron, and fast-spiking interneuron dynamics. All the parameter values and expressions here described were taken from (Rotstein, et al., 2005).

| Parameter | Value |
| --- | --- |
| **O-cells** |  |
| $C_m$ | $1 \mu F/cm^2$ |
| $g_{leak}$ | $0.5 mS/cm^2$ |
| $E_{leak}$ | $-65 mV$ |
| $\bar{g}_K$ | $11 mS/cm^2$ |
| $E_K$ | $-90 mV$ |
| $\bar{g}_{Na}$ | $52 mS/cm^2$ |
| $E_{Na}$ | $55 mV$ |
| $\bar{g}_p$ | $0.5 mS/cm^2$ |
| $\bar{g}_h$ | $1.45 mS/cm^2$ |
| $E_h$ | $-20 mV$ |
| **I-cells** |  |
| $C_m$ | $1 \mu F/cm^2$ |
| $g_{leak}$ | $0.1 mS/cm^2$ |
| $E_{leak}$ | $-66 mV$ |
| $\bar{g}_K$ | $80 mS/cm^2$ |
| $E_K$ | $-100 mV$ |
| $\bar{g}_{Na}$ | $100 mS/cm^2$ |
| $E_{Na}$ | $50 mV$ |

Since we are interested in studying local synaptic changes of the SC-CA1 synapse, we use the following equation to describe the activity of the pyramidal cell dendritic compartment:

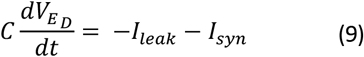

The parameters C, g_leak_ and E_leak_ are the same as for the I-cells.

### Synaptic models

The O-cells have a synaptic current mediated by pre-synaptic α7 nAChR channels. The description of the current here used is an adaptation of the model proposed in (Graupner & Gutkin, 2013), and it is given by

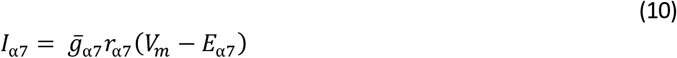

where 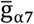 is the maximal conductance of the α7 nAChR channel, and *E*_α7_ the reversal potential. The opening gate variable r_α7_ is described by equation (8), with τ_rα7_ constant and r_(α_7_)∞_ given by

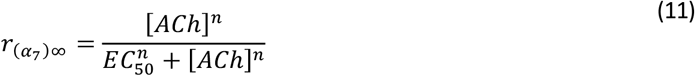

where EC_50_ is the half-maximum concentration, and *n* the Hill’s coefficient of activation.

The I-cell has an excitatory AMPA and inhibitory GABA_A_ synaptic currents, described by the following set of equations:

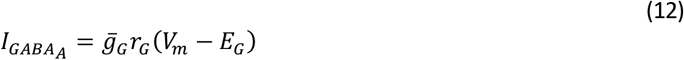

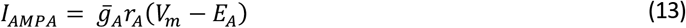

The gating variables *r_x_* is, as described in (Destexhe, Mainen, & Sejnowski, 1998), given by

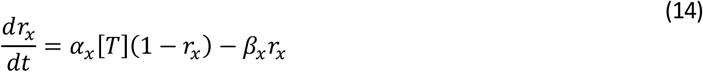

where α_x_ and β_x_ are the opening and closing rate of the receptor channel, and [T] the neurotransmitter’s concentration available for binding.

The GABA release resultant from the respective activation of the O and I-cells depends on its membrane potential V_m_ according to the following equation:

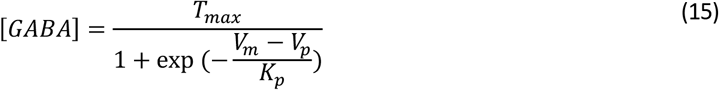

where *T_max_* is the maximal neurotransmitter concentration, *K_p_* gives the steepness of the function, and *V_p_* sets the value at which the function is half-activated.

The dendritic compartment E_D_ is modeled using synaptic GABA_A_, AMPA and NMDA currents. The GABA_A_ and AMPA currents are given by equations (12) and (13), respectively. The NMDA current is described by the following equation:

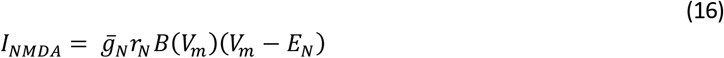

where *r_N_* is the gating variable described by equation (14). Due to the presence of a Mg^2+^ block, the NMDA channels have an extra voltage-dependent term, B(V_m_), defined as:

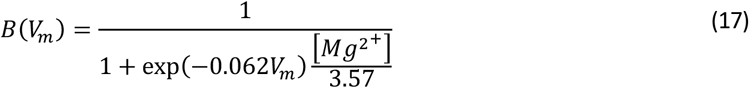

The reversal potentials of the excitatory receptors *E_A_* and *E_N_* are 0 mV, while one of the inhibitory GABA_A_ receptor is E_G_ = −80 mV. Postsynaptic AMPA and GABA_A_ receptors at the I-cell have maximal conductance 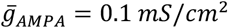 and 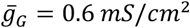. at *E_D_* have a maximal conductance of 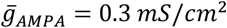. Postsynaptic AMPA, GABA_A_ and NMDA receptors at *E_D_* have maximal conductances of 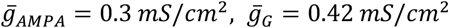 and 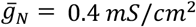, except for Figures 2 and 3 where we use 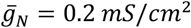 and 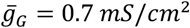. The remaining parameters are defined in Table 2.

**Table 2:**
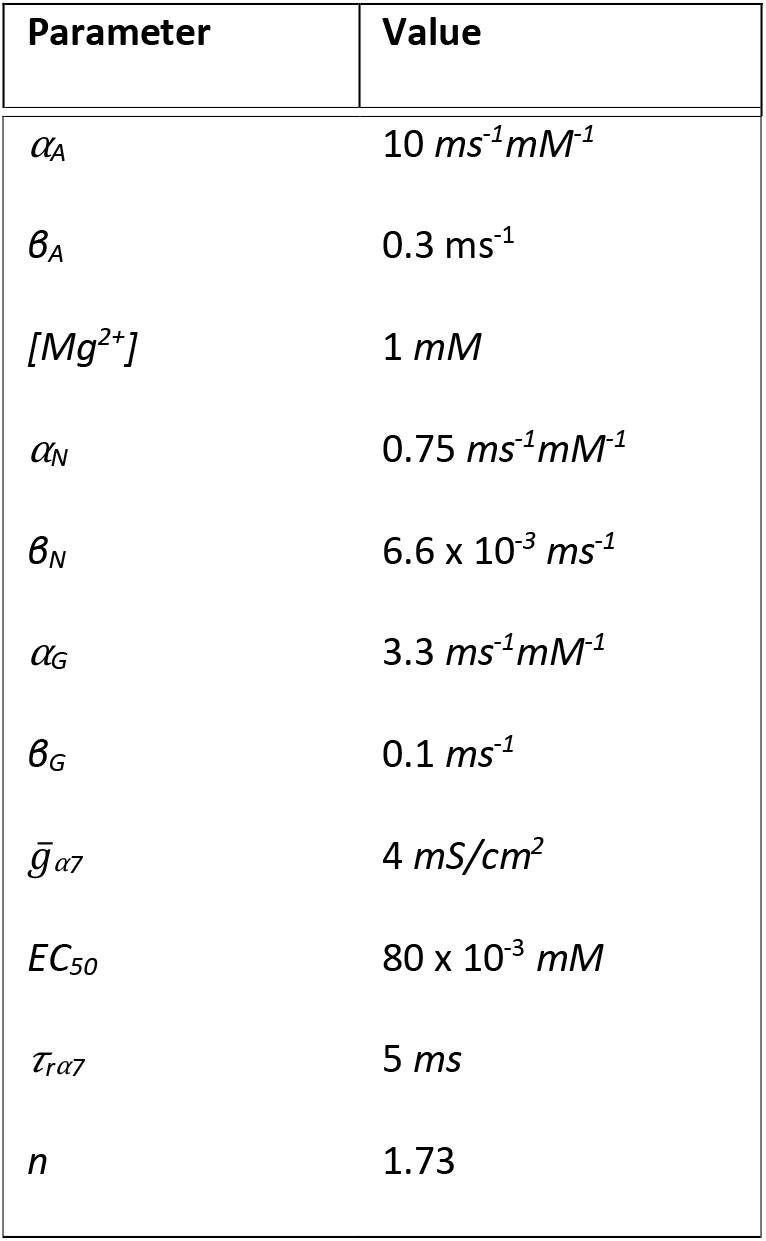
Parameter values of synaptic currents *AMPA, NMDA* and *GABA_A_*. The parameter values were based on (Graupner & Gutkin, 2013) and (Destexhe, Mainen, & Sejnowski, 1998).

### Model of synaptic plasticity

We use a calcium-based synaptic plasticity model based on (Shouval, Bear, & Cooper, 2002). We assume that changes in the AMPA receptor conductance reflect changes in the strength of the excitatory SC-CA1 synapse. Our synaptic plasticity model is formulated as follows:

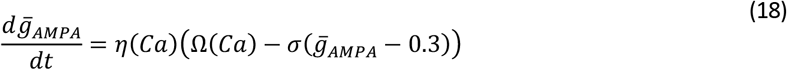

where *σ* is a decay constant. The variable *η* is a calcium-dependent learning rate described by equation (19), and Ω determines the sign magnitude of synaptic plasticity as a function of the intracellular Ca levels (equation (20)).

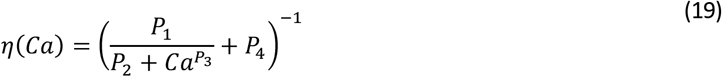

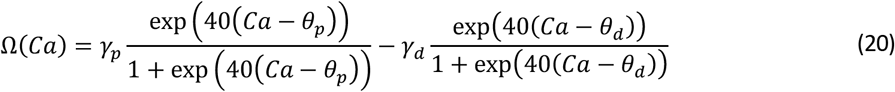

The parameters Θ_*p*_ and Θ_*d*_ define the depression and potentiation threshold, with Θ_*p*_ > Θ_*d*_, and *γ_p_* and *γ_d_* define the potentiation and depression amplitude, with *γ_p_* > *γ_d_*.

We assume that the main source of *Ca*^2+^ is the calcium flux entering the cell through the NMDA receptor channels. The intracellular *Ca*^2+^ concentration evolves according to the following equation

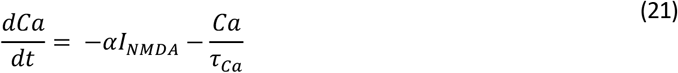

where *α* is the NMDARs permeability to calcium, and *Tea* the calcium decay constant.

The parameter values used in the simulations are the ones presented in Table 3, except for Figure 2 and Figure 3 where *γ_p_* = 0.165 and *γ_d_* = 0.100.

**Table 3:** Parameter values for synaptic plasticity. The values were adjusted from (Shouval, Bear, & Cooper, 2002) to obtain calcium level changes able to induce potentiation and depression in our simulations.

| Parameter | Value |
| --- | --- |
| $\sigma$ | 0.003 |
| $P_1$ | 30 *ms* |
| $P_2$ | $P_1 \times 10^{-4}$ |
| $P_3$ | 11.4 |
| $P_4$ | 1 *ms* |
| $\vartheta_p$ | 1.4 |
| $\vartheta_d$ | 1 |
| $\gamma_p$ | 0.143 |
| $\gamma_d$ | 0.105 |
| $\alpha$ | 0.2 |
| $\tau_{Ca}$ | 5 *ms* |

## Results

### Co-activation of cholinergic and glutamatergic inputs modifies the SC-CA1 synaptic transmission

Previously, it has been observed that coactivation of hippocampal cholinergic inputs and local SC pathway increases the amplitude of SC to CA1 pyramidal EPSCs. Moreover, repeated pairing of cholinergic and hippocampal inputs (8 times) induced the enhancement of EPSCs from pyramidal neurons not only during the pairing, but also long after, indicating the induction of long-term synaptic plasticity at SC to CA1 excitatory synapses. The induction of potentiation was abolished by a knock-out of OLMα2 interneuronal α7 nAChRs (Figures 2A and 2C).

SC stimulation elicits EPSCs in s.r. interneurons and in the CA1 pyramidal cell dendrites (Leão, et al., 2012). On the other hand, despite the fact that α7 nAChR-mediated currents have been observed in both hippocampal pyramidal neurons and interneurons, only the knockout of α7 nAChRs in OLMα2 interneurons blocked the potentiation of cholinergic-pairing-induced SC-evoked pyramidal cell EPSPs (Gu, et al., 2020). We thus modeled α7 nAChR-mediated modulation of OLM neurons only. We constructed a minimal feed-forward circuit with an OLM cell (O), a fast-spiking interneuron (I), and the pyramidal cell s.r. dendritic compartment (E_D_) connected as schematically shown in Figure 1B to examine mechanistically how pairing cholinergic activation of the O-cell with glutamatergic activation of the I-cell and E_D_ can potentiate the EPSCs of E_D_. Both the OLM cell and the FS neuron are modelled following a conductance-based Hodgkin-Huxley formalism (see methods). We study how the EPSC of E_D_, modelled as the sum of the postsynaptic AMPA and NMDA currents (I_AMPA_ and I_NMDA_), change when the glutamatergic inputs acting on the I-cell and E_D_ are paired with the cholinergic inputs that act on the post-synaptic α7 nAChR of to the O-cell during a co-pairing period of 8 minutes, identical to the experimental protocol. Both before, during and after the co-pairing period, the I-cell and E_D_ receive one pulse of glutamate per minute. During the co-pairing period the O-cell receives a pulse of ACh per minute, 1 msec prior to each glutamate pulse. Not much is known about the concentration profile of ACh in vivo, but it is believed that it can be cleared from the synaptic cleft within milliseconds. Therefore, both ACh and glutamate are modeled as square pulses with a duration of 3 msec and 1 mM of amplitude. ACh then acts to activate the α7 nAChRs, producing a depolarising current in the target neuron in our model.

From Figure 2D, we see that during the co-pairing period (from t=10 min to t=18 min), the EPSC is increased. This increase in our model is maintained for an extended period of time after the co-pairing period is over (black line), matching the experimental results. We also see that during the co-pairing period, GABA release from the I-cells, *GABA_I_*, decreases significantly (see Figure 2D inset). If we decrease the maximal conductance of the a7 nAChR, *g_α7_*, from 4 mS/cm^2^ to 1 mS/cm^2^ (or lower), co-pairing no longer potentiates the EPSC of E_D_ (orange line). This is in accordance with experimental results that showed that this form of EPSC boost was abolished by knockout of the α7 nAChR in OLMα2 interneurons (Figure 2C).

We then examined how the key parameters of the co-paring protocol influence the plasticity of the SC-CA1 EPSCs. According to our model, the duration of the co-pairing period, the relative time between the cholinergic and glutamatergic inputs, as well as their frequency during the co-pairing period, can modulate the efficiency and direction of plasticity. Results of our simulations show that the longer the co-pairing period, the longer will be the transient duration, where the potentiation transient duration was defined as the time it takes the EPSCs to return to its baseline value once the co-pairing period is over (Figure 3A). We observe a positive relationship between the frequency of the glutamatergic and cholinergic inputs during a fixed period of paring protocol, and the duration of the potentiation transient (Figure 3B). Interestingly, our simulations suggested that while changing the co-pairing period and the frequency of stimulation modulates the efficiency of the induction of potentiation, it does not change the direction of plasticity. It was only when varying the relative time between the ACh and glutamate pulses that we could induce a change in the direction of the plasticity. For the pairing of single pulses, if the glutamatergic inputs arrive to I and E_D_ within 0.7 < Δ*t* < 20.2 msec following the ACh pulse potentiation will be induced. If 20.2 < Δ*t* < 38 msec, depression is induced (Figure 3C). If instead of single pulses we pair doublets of glutamate and ACh, the potentiation and depression window change. The potentiation window is of 0.3 < Δ*t* < 18.27 msec, while the depression window is −6.28 < Δ*t* < 0.3 msec and 18.27 < Δ*t* < 70 msec (see Figure 3D and S1).

### Disinhibition of the CA1 pyramidal cell dendritic compartment enables potentiation of the SC-CA1 synaptic transmission

As we have shown in Figure 2, our model shows a decrease in GABA release from I-cells during the co-pairing period. Based on this result, we hypothesize that a decrease of GABAergic inhibition of E_D_ allows for the potentiation of the SC-evoked EPSC of the pyramidal cell dendritic compartment. To study the role of disinhibition of E_D_ in the potentiation of the SC-CA1 excitatory synapse, we use a model where E_D_ receives a pulse of glutamate followed by a pulse of GABA, except during a disinhibition period when it only receives pulses of glutamate. We also paired, *in vitro*, inhibition of Sst interneurons with SC stimulation.

From Figure S2 we see that the rise and decay time of GABA concentration release that results from the spiking of the I-cells is almost instantaneous. Therefore for simplification, in this section GABA is modelled as a square pulse. Both glutamate and GABA release pulses are modeled as square pulses with a duration of 1 msec and 1 mM of amplitude. E_D_ receives one pulse of glutamate per minute, followed by a pulse of GABA 2 msec after, except during a disinhibition period when it only receives pulses of glutamate. We note that this choice of simulated stimulation and pairing directly follows from the experimental protocol (see Methods). We use such a model to study how can disinhibition of E_D_ modulate the induction of synaptic plasticity.

It was observed that during the disinhibition period the EPSC amplitude increases, and that the longer the disinhibition period last, the longer these changes last. Specifically, our model shows that for a disinhibition period of 4 minutes, the EPSC returns to baseline once the disinhibition period is over, and that for a longer disinhibition period of 8 minutes, the EPSC remains potentiated long after the disinhibition period is over (Figure 4D). This is in accordance with experimental results, where inhibition of Sst interneurons projecting to CA1 pyramidal cells was paired with SC stimulation for a short and long period of time (Figure 4C). Inhibition of Sst interneurons via eNpHR resulted in increased SC-CA1 EPSC amplitude not only during the Sst inhibition, but also after the end of Sst inhibition. The EPSC enhancement after the end of Sst inhibition lasted about 10min after 4 times of Sst inhibition and more than 30 min after 8 times of Sst inhibition.

Given that the amplitude of the AMPA current is much larger than the NMDA current (a glutamate pulse with a duration of 1 msec and 1 mM of amplitude elicits an AMPA current with an amplitude of 17.65 *μ*A/cm^2^ and a NMDA current with an amplitude of 2.19 *μ*A/cm^2^), the AMPAR arethe greatest contributor for the EPSC of E_D_. Therefore, we deduce that the changes in the EPSC amplitude seen during the disinhibition period reflect changes in the AMPARs maximal conductance, 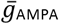. In fact, AMPARs have been known to play an important role in the regulation and expression of synaptic plasticity in the hippocampus (Barria, Muller, Derkach, Griffith, & Soderling, 1997). From Figure 5 we see that there is an increase of 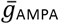 during the disinhibition period. The longer the disinhibition period, the higher is the value of 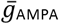 when the disinhibition period is over. For a disinhibition period of 4 minutes (blue line), there is an increase of 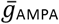 from 0.3 to 1.8 *mS/cm*^2^ during disinhibition. Afterwards, 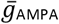 gradually goes back to its baseline value. For a disinhibition period of 8 minutes (orange line), 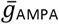 increases from 0.3 to 2.8 *mS/cm*^2^. When the disinhibition period is over, 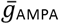 remains potentiated.

**Figure 5:**
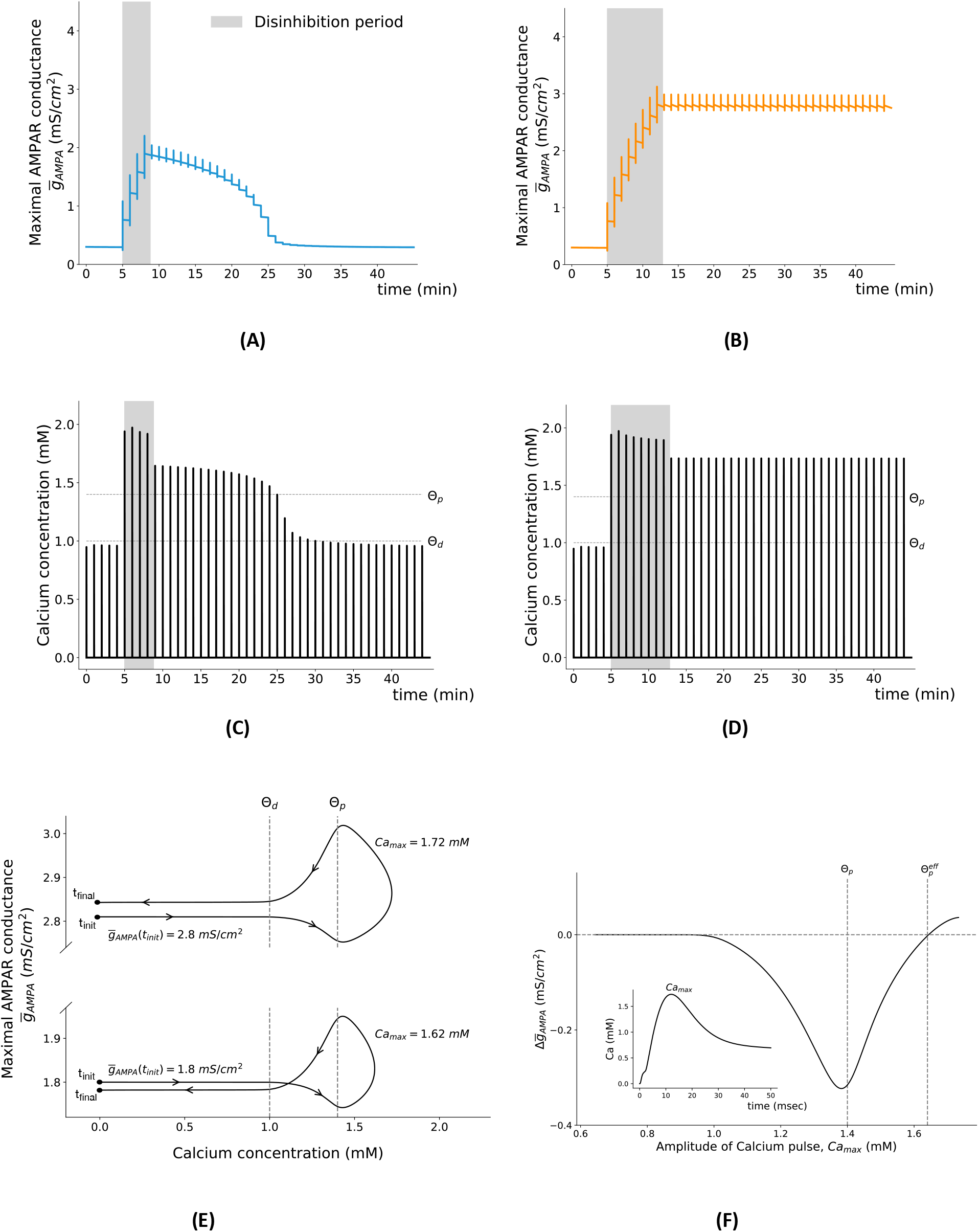
Calcium dynamic is key for the induction of synaptic plasticity. **(A)** Time course of maximal AMPAR conductance, 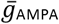, when dendritic compartment is disinhibited for a short period of time (from t=5 min to t=9 min). The maximal AMPAR conductance increases from its initial value 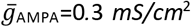 to 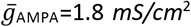 during disinhibition period. **(B)** Time course of 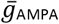 when dendritic compartment is disinhibited for a long period of time (from t=5 min to t=13 min). It increases from its initial value 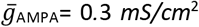 to 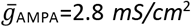 during the disinhibition period. The AMPAR conductance 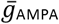 is described by equation 18. **(C)** Time course of intracellular calcium concentration when dendritic compartment E_D_ is disinhibited for a short period of time (from t=5 min to t=9 min). The pairing of a pulse of glutamate with a pulse of GABA at t=10 min, i.e. immediately after the disinhibition period, results in a calcium pulse with an amplitude of 1.62 mM. **(D)** Time course of intracellular calcium concentration when dendritic compartment is disinhibited for a long period of time (from t=5 min to t=13 min). The pairing of a pulse of glutamate with a pulse of GABA at t=14 min, i.e. immediately after disinhibition period, results in a calcium pulse with an amplitude of 1.72 mM. The calcium dynamics is described by equation 21 (see Methods). **(E)** Trajectories of the system in the 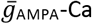 plane when a pulse of glutamate is paired with a pulse of GABA for different initial values of 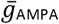. **(F)** Changes in the maximal AMPAR conductance, 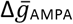, as a function of the amplitude of intracellular calcium pulse, Ca_max_. Each point of the graph was obtained by submitting E_D_ to a pulse of glutamate for different initial values of 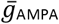. This induced different levels of depolarization and, consequently, different levels of activation of NMDARs and calcium pulses of different amplitudes.

In our simulations we saw hints that not only the maximal voltage-dependent calcium concentration crossing the potentiation threshold set in the model, but also the dynamics of calcium are determinant for the synaptic plasticity. Hence, we examined the dynamics of the intracellular calcium concentration to understand how it determines disinhibition-driven synaptic plasticity. From Figures 5C and D we see that calcium concentration increases significantly during the disinhibition period, crossing the potentiation threshold Θ_p_ with a significant margin. For both the short and long disinhibition, after the end of the disinhibition period, the calcium levels decrease, yet they remain above Θ_p_. This clearly shows that the maximal calcium concentration, Ca_max_, is not the sole determinant of long-term synaptic potentiation. In the case of a short disinhibition period, each pairing of GABA and glutamate after the disinhibition period will elicit a calcium pulse with a smaller amplitude than the previous one. Eventually, at t=25 min, the calcium concentration that results from the pairing is not enough to cross the potentiation threshold, and by the time t=30 min it does not cross either the potentiation or the depression threshold, having a similar amplitude as before the disinhibition period. In the case of a long disinhibition period, each pairing performed after the disinhibition period evokes a calcium pulse with the same amplitude. To better visualize the synaptic and calcium dynamics immediately after the disinhibition period in both cases, we plot the system’s trajectory in the 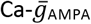 plane. We do so for 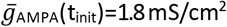 and for 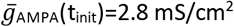 (Figure 5E), which are the values of 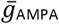 at the end of the disinhibition period for the short- and long-disinhibition durations. For 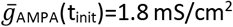, the calcium concentration crosses the potentiation threshold Θ_p_ (Ca_max_ = 1.62 mM) but there is a total decrease in 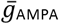. For 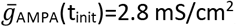, the calcium concentration crosses Θ_p_ to a larger extent (Ca_max_ = 1.72 mM) and there is a total increase of 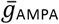. This result reveals the importance of calcium dynamics in the induction of plasticity. Changing the calcium dynamics, for example by changing the decay constant of calcium, will change the outcome of plasticity induction by changing the time calcium spends just above the potentiation threshold – when potentiation is induced – and between the depression and potentiation threshold – when depression is induced (see Fig. S4).

These results also suggest that it is not sufficient for calcium concentration to cross the potentiation threshold to result in a potentiation of 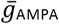. To verify this, we looked at changes in maximal conductanceof the postsynaptic AMPAR, 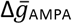, as a function of the amplitude of the intracellular calcium, Ca_max_. From Figure 5F we see that as Camax increases we only start to have potentiation 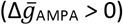 when Ca_max_ crosses not the potentiation threshold Θ_p_, but what we will call an effective potentiation threshold Θ^eff^, at Ca_max_=1.61 mM.

Based on our findings, we infer that the disinhibition period can govern the efficiency of the plasticity induction by controlling the extent to which 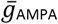 is potentiated, which in turn will determine the outcome of the stimulation protocol after disinhibition period.

Similarly to what was done in the previous section, we wanted to examine how other parameters influence the plasticity of SC-CA1 EPSCs. According to our model, the amplitude of the GABA pulse, GABAmax, as well as the relative time between the glutamate and GABA pulses, Δt_(GABA-Glu)_, can modulate plasticity. We also see that the regions of the Δt_GABA-Glu_-GABA_max_ parameter space that define the zones of depression and potentiation changes with the 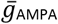 (Figure 6). Therefore, the same induction protocol may induce potentiation or depression, more or less efficiently, depending on the current state of phosphorylation of the AMPA receptors, i.e. 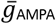, and on the decrease of GABA during disinhibition.

**Figure 6:**
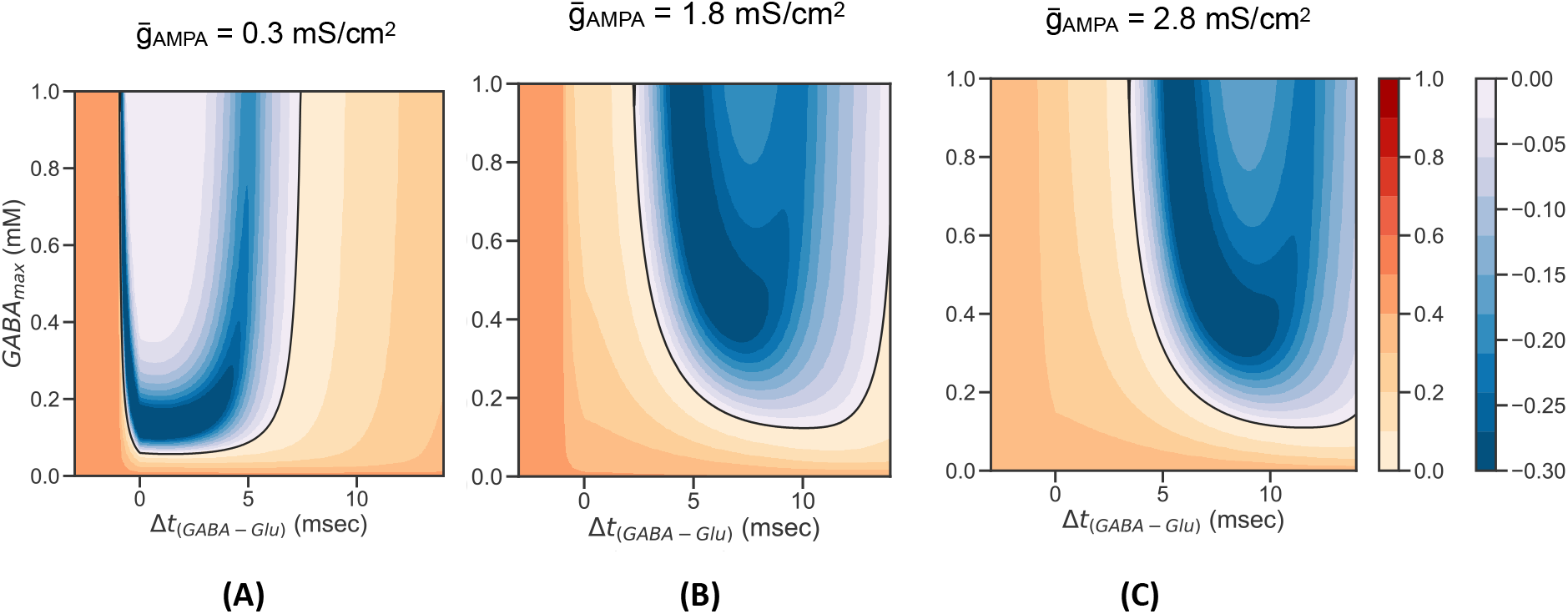
Amplitude of GABA pulse, GABA_max_, and relative time between GABA and glutamate pulses, Δ*t*(*GABA-Glu*), control direction and efficiency of induction of synaptic plasticity. **(a)** Depression and potentiation regions for 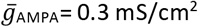. **(b)** Depression and potentiation regions for 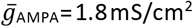. **(c)** Depression and potentiation regions for 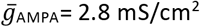.

### Membrane potential of CA1 pyramidal dendritic compartment controls direction of plasticity

Based on our results, we hypothesize that the voltage of the pyramidal dendritic compartment V_E_D__ is the critical variable in the model that controls the direction of plasticity. To explore this hypothesis, we will use phase-plane analysis. To use this technique, we reduce our 6-variable model (V_E_D__, 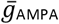, r_N_, r_G_, r_AMPA_, Ca) to its slow 2-variable subsystem (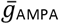, r_N_) so that it may be represented graphically and understood entirely by doing phase-plane analysis. We do this by exploring the time scale differences between the dynamical variables of the system. The maximal conductance of the AMPAR and the fraction of open NMDA receptor channels, 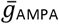 and *r_N_*, are significantly slower than the other variables. We use a quasi-equilibrium approximation to the fast variables Ca, r_AMPA_ and r_G_, i.e. we replace this variables by their steady state value, to obtain a reduced 2-variable 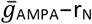 with V_E_D__ as a parameter. We then plot the trajectories and respective nullclines of the reduced system in the 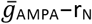 plane for 2 different values of V_E_D__, V_E_D__ = −69 mV and V_E_D__ = −60 mV (Figure 7). In the reduced 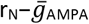 system, r_N_ is the fast variable, while 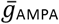 is the slow variable. We note that from the dynamical systems theory we know that in such fast-slow systems, the solutions tend to move rapidly towards (or away from) the fast variable nullclines, and then flow slowly along it (the so-called slow manifold). Hence for most of the time, the dynamics of the system are functionally governed by the shape and the location of the fast nullcline relative to the slow one and the slow-variable flow along it. More precisely, starting from some general initial condition, the system will be governed by the fast subsystem rN until it reaches its steady state 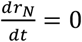. The system will then evolve slowly governed by the slow subsystem 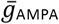. As shown in Figure 7, the parameter V_E_D__ controls the nullcline of 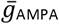. For V_E_D__ = −69 mV, the nullcline of rN is below the one of 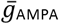. Therefore, when the steady state of r_N_ is reached, the system will be driven to the left, which corresponds to a decrease in 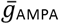 - or, in other words, to depression. On the other hand, when V_E_D__ is increased to −60 mV the nullcline of rN is now located above 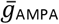, and the system will evolve towards the right once the steady state of rN is reached - we have potentiation.

**Figure 7:**
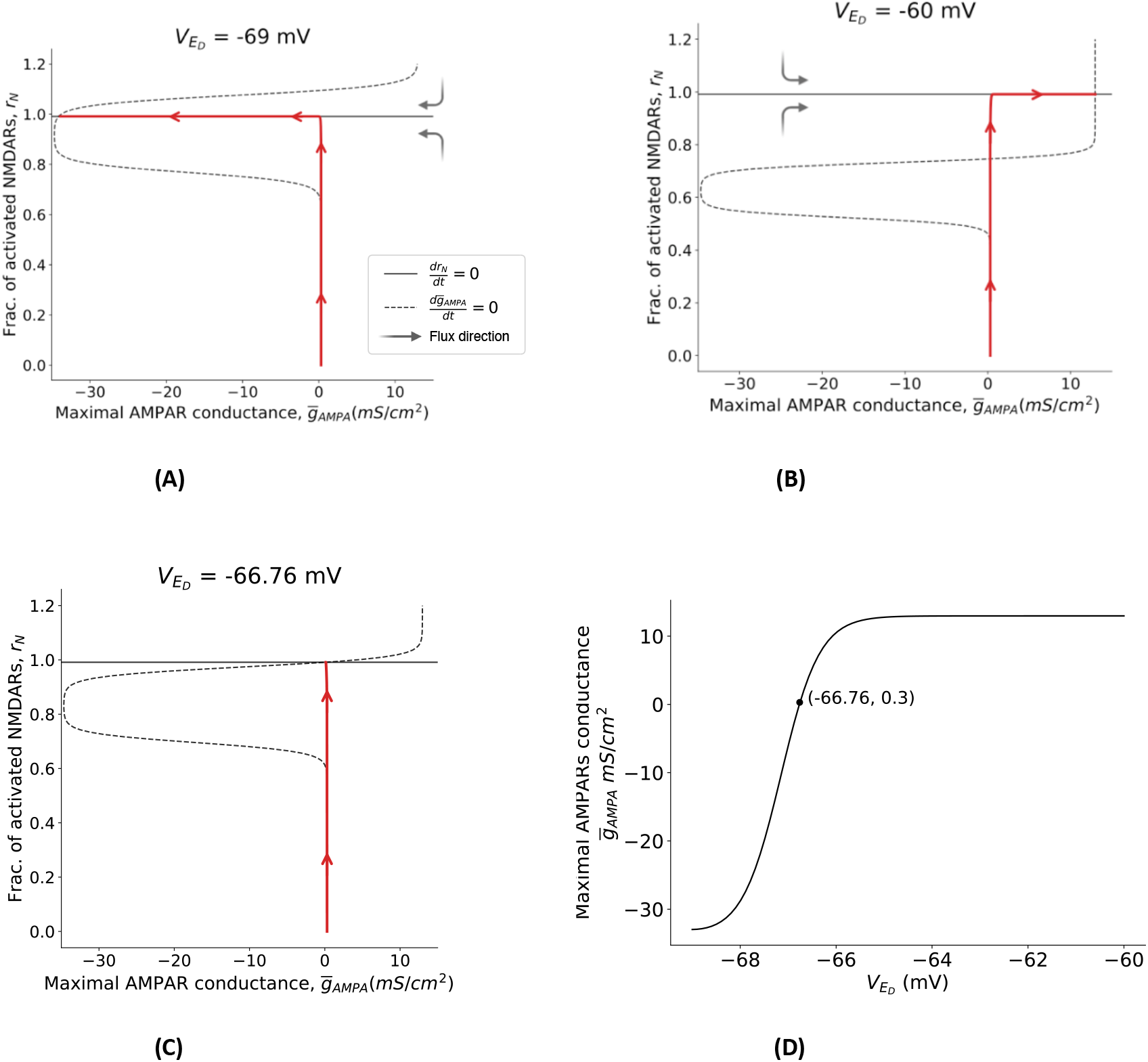
**Trajectories of the slow subsystem 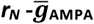 for V_eD_ =-69 mV and V_eD_ =-60 mV, and respective nullclines.** **(A)** For V_eD_ = −69 mV, the system has a stable fixed point at (−34.1,0.99). Initiating the system at 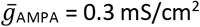 and r_N_ =0, depression will be induced. **(B)** For V_eD_ = −60 mV, the system has a stable fixed point at (12.9,0.99). Initiating the system at 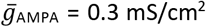 and r_N_ =0, potentiation will be induced. **(C)** For V_eD_ = −66.76 mV, the system has a stable fixed point at (0.3, 0.99). Initiating the system at 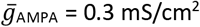 and r_N_ =0, no plasticity will be induced. **(D)** Bifurcation diagram for slow subsystem 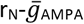. The system has a continuity of stable steady states. If we initialize the system with 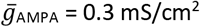, values of V_ed_ bellow −66.76 mV will move the system to a steady state with 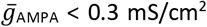, inducing depression. Values of V_ED_ above −66.76 mV will make the system evolve towards a steady state with 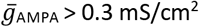, inducing potentiation.

These results suggest that a necessary condition for the induction of synaptic plasticity is the depolarization of the pyramidal cell membrane, which can be accomplished by pairing the activation of the SC with the disinhibition.

### Model Predictions

Results of model simulations and analysis make several testable predictions. Despite not knowing the exact type of s.o. interneuron that is providing feedforward inhibition to the CA1 pyramidal cell, our model predicts that it should be an interneuron with fast dynamics, i.e. with dynamics comparable to the one of the pyramidal cells.

In this work (both in modelling and experimentally), activation of OLM cells is accomplished through cholinergic activation of α7 nAChRs. In our model, α7 receptor activation provides a depolarising current to the OLM neurons. Therefore, we expect that activation of the OLMα2 interneurons through any mechanism other than septal cholinergic activation, for example through optogenetic stimulation, would lead to the similar results.

Finally, our model predicts a relationship between the relative timing of the septal and hippocampal stimulus paring and the direction of the synaptic plasticity at the SC-PYR synapse. According to our simulations, increasing the frequency of septal and hippocampal paired stimulation can induce plasticity more easily, i.e. less pairings would be required to induce LTP. At the same time, we predict that increasing the relative time between septal and hippocampal activation can induce LTD instead of LTP.

## Discussion

This work set out to explain how cholinergic activation of hippocampal OLM interneurons paired with hippocampal stimulation can potentiate CA1 pyramidal cell EPSC responses. Our modelling results suggest that co-pairing cholinergic activation of OLM interneurons results in the disinhibition of CA1 pyramidal cells, which paired with hippocampal stimulation potentiates CA1 pyramidal cell SC-evoked EPSCs. We also show general proof of principle of how synaptic plasticity is controlled by the disinhibition of the postsynaptic pyramidal membrane through a disynaptic GABA-ergic circuit.

OLM cells are a major class of GABAergic interneurons located in the stratum oriens hippocampal layer that inhibit pyramidal cells dendritic compartment located in the stratum lacunusom-moleculare layer, reducing the strength of EC inputs. However, there are recent findings showing that activation of these cells can facilitate LTP in SC-CA1 pathway, likely by inhibiting s.r. interneurons that synapse on the same dendritic compartment as the SC, counteracting SC feedforward inhibition (Leão, et al., 2012).

We found that repeated pairing of cholinergic inputs with hippocampal stimulation can induce plasticity if the inputs are tightly timed. The time window for potentiation depends greatly on the dynamics of the O-cells, I-cells and the calcium dynamics. This is in accordance with experimental findings showing that activation of cholinergic inputs to the hippocampus can directly induce different forms of synaptic plasticity depending on the input context in the hippocampus, with a timing precision in the millisecond range (Gu & Yakel, Timing-dependent septal cholinergic induction of dynamic hippocampal synaptic plasticity, 2011). Our model also shows that the longer the co-pairing period and the higher the frequency of stimulation during the co-pairing period, the longer lasting is the potentiation of the synapse.

It is important to note that in our model, the OLM fast-spiking interneuron and pyramidal cell are connected through a feedforward connection. It is likely that recurrent connections between these neurons exist. However, adding these recurrent connections to our model does not qualitatively change our results, provided we adjust the parameters corresponding to the potentiation and depression threshold, Θ_p_ and Θ_d_, and/or the potentiation and depression amplitudes, γ_p_ and γ_d_, as well as the maximal conductance of the synaptic currents.

According to our model, the key mechanism behind paired cholinergic induction of synaptic plasticity is the disinhibition of the pyramidal cell dendritic compartment. Cholinergic activation of the O-cell inhibits the fast-spiking I-cell that projects to the dendritic compartment E_D_. The disinhibition of E_D_ paired with glutamatergic stimulation allows for the depolarization of the pyramidal dendritic compartment. This leads to increased NMDAR activation and intracellular calcium concentration sufficient to upregulate postsynaptic AMPAR permeability and potentiate the excitatory synapse. It is worth mentioning that we consider a minimal model and that there are other pathways that may play a role in the induction of plasticity by modulating the calcium dynamics at the s.r. dendritic compartment; namely, the OLM projection to the distal dendrites of the CA1 pyramidal neurons.

In order to study in greater detail the mechanisms behind the induction of plasticity through disinhibition, we modelled changes in synaptic strength when glutamatergic activation of E_D_ is paired with locally reduced GABA.

Before the disinhibition period starts, the cell receives both excitatory and inhibitory inputs. An excitatory input evokes an EPSP mediated by AMPAR. The EPSP is followed by a GABA_A_-mediated IPSP. Because of slow kinetics and voltage-dependence, at that time, the NMDAR receptors are not in the open state and there is no influx of calcium. During the disinhibition period, the cell only receives the excitatory inputs (GABA inputs are suppressed). The pyramidal cell membrane at (or sufficiently near to) the glutamatergic synapse is able to depolarize enough to relieve the Mg^2+^ block from the NMDA receptors, allowing calcium to permeate through the receptor channel (Figure 8). Therefore, during the disinhibition period, every time the pyramidal cell receives a pulse of glutamate, the intracellular calcium concentration crosses the potentiation threshold Θ*p*, and 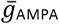 increases.

**Figure 8:**
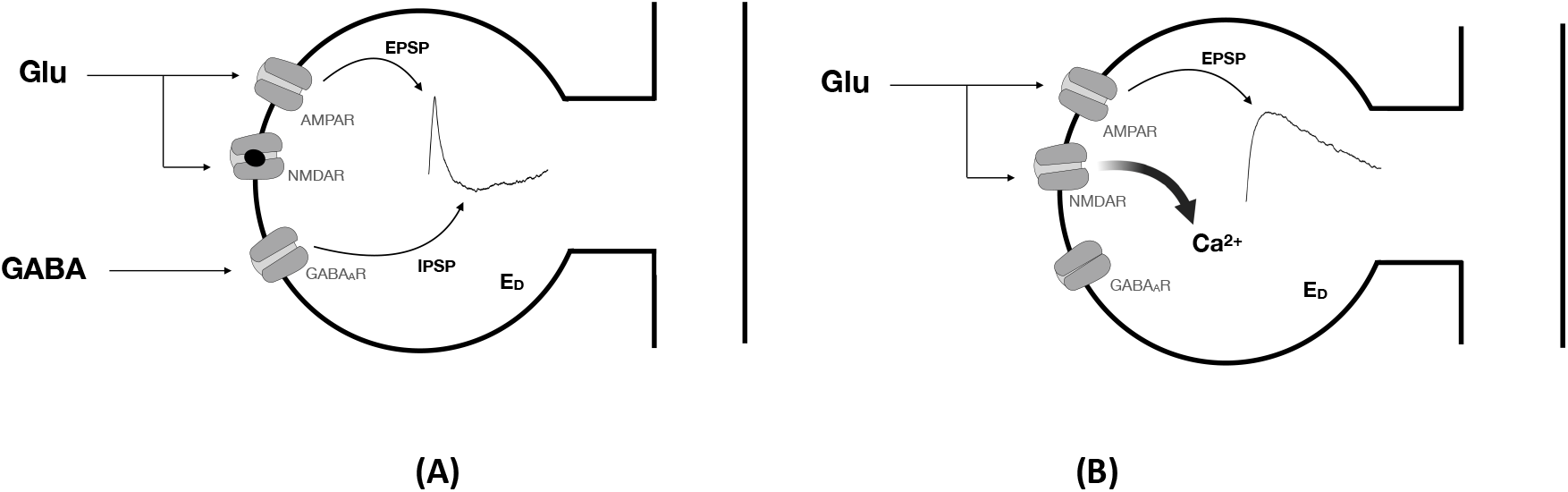
Scheme of the activation mechanisms of NMDARs and consequent calcium influx. **(A)** Glutamatergic activation evokes an EPSP mediated by AMPAR. The EPSP is followed by an IPSP mediated GABA acting on GABAA receptors. **(B)** Dendritic compartment does not receive GABAergic inhibition. The dendritic compartment is able to depolarize enough - and remain depolarized for long enough - to relieve Mg^2+^ block from NMDA receptors, allowing calcium to permeate through the receptor channel.

Therefore, down-regulation of the GABAergic signaling during disinhibition leads to increased NMDAR activation. Because of the receptor’s high calcium permeability, there is an elevation in intracellular calcium concentration large enough to initiate molecular mechanisms that result in the insertion/phosphorylation of the AMPAR. In our model, this translates into an increase in the AMPAR maximal conductance 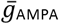. Moderate calcium concentrations, on the other hand, result in the removal of AMPARs. Because changes in calcium concentration are not instantaneous, the potentiation/depression of the synapse is a result of a balance between the insertion/removal of AMPARs during the period in which Ca concentration is above potentiation/depression threshold. During disinhibition, this balance is positive and there is a total increase in 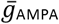. The more times we pair disinhibition with SC stimulation, i.e. the longer the disinhibition period, the higher will be the value of 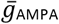 by the end of the disinhibition period. After the disinhibition period, if the increase of 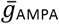 was large enough, the calcium resultant from glutamatergic and GABAergic stimulation is such that there was a balance between potentiation and depression, which is close to zero 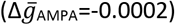. Therefore, the synapse remains potentiated long after the disinhibition period is over. If there is no stimulation after the disinhibition period, 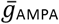 slowly decays to its initial value (i.e. its value before the disinhibition period). If the increase of the AMPAR permeability is high enough, the potentiation of the excitatory synapse is maintained when the disinhibition period is over through repeated stimulation of the SC that keeps a balance between the down and upregulation of the AMPARs. This is in accordance with experimental results that show that repeated pairing of inhibition of Sst interneurons (which are not OLM interneurons) that target the CA1 pyramidal cell with SC stimulation can induce plasticity.

Our model is robust to changes of parameters that maintain the same ratio of potentiation/depression. Meaning that, for example, for different values of the potentiation amplitude Yp, there is (at least) a pair of maximal depression Yd and decay constant *σ* for which our results remain the same (Fig. S3).

It is worth noting that the type of synaptic plasticity induced depends on the value of maximal conductance of the postsynaptic AMPAR, 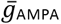, as shown in Figure 6. Therefore, despite not having explicit memory or considering metaplasticity, our model shows that future changes in synaptic strength depend on previous plasticity events and how these changed 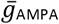. This explains why, after the disinhibition period, pairs of glutamate and GABA pulses with the same characteristics are going to induce different results when the disinhibition period was short or long.

In summary, using our minimal biophysically-based model, we accounted for key experimental observations that provided a cogent explanation for the role of the septal cholinergic pathway activation in the induction of hippocampal synaptic plasticity. Our modeling work suggests that GABAergic neurotransmission controls the local pyramidal voltage in the vicinity of the glutamatergic synapses, thereby the inhibitory synapses critically modulate excitatory transmission and the induction of plasticity at excitatory synapses. This puts into evidence the importance of dendritic GABA and glutamate co-location in shaping local plasticity rules. Our work also suggests a cholinergic mechanism for the inhibition of GABA release at the pyramidal dendrites and consequent potentiation of excitatory synapses, unraveling the intricate interplay of the hierarchal inhibitory circuitry and cholinergic neuromodulation as a mechanism for hippocampal plasticity.

Previous work by Gu et al. (2017) showed that co-paired activation of the cholinergic input pathway from the septum to the hippocampus, either electrically or using optogenetics, with stimulation of the Schaffer collateral pathway could readily induce theta oscillations in a co-culture septal-hippocampal-entorhinal preparation. They also observed that performing this co-pairing mechanism increased excitatory transmission in the hippocampus and subsequent outputs to the EC in slice co-cultures. Moreover, after synaptic plasticity has been induced it becomes easier to evoke the theta rhythm in the preparation (one pulse stimulus of the SC is sufficient to generate theta oscillations in the circuit with the same characteristics as before) (Gu & Yakel, Inducing theta oscillations in the entorhinal hippocampal network in vitro, 2017; Gu, Alexander, Dudek, & Yakel, 2017). Therefore, induction of hippocampal plasticity, in particular potentiation of the CA1 EPSPs, appears to facilitate the generation of the theta rhythm. Moreover, a recent study directly linked OLMa2 interneurons to theta oscillations (Mikulovic, et al., 2018). We thus may speculate that the action of ACh on the a7 nAChRs at the OLMa2 neurons potentiates the SC-CA1 synapses to close the critical link in the synaptic chain of events, enabling recurrent reverberation of excitation in the hippocampal-entorhinal theta generating circuit. Understanding the mechanisms underlying the induction of hippocampal plasticity by this co-pairing mechanism will allow future studies of how changes on the synaptic level can propagate to the network level and change the mechanisms of theta generation.

We may also note that our results can be relevant to understanding the neural circuit origins of pathological conditions. The hippocampus is one of the earliest brain structures to develop neurodegenerative changes in Alzheimer’s disease (AD) (Arriagada, Growdon, Hedley-Whyte, & Hyman, 1992). Numerous studies suggest that cognitive deficits in AD such as memory impairment are caused in part by cholinergic dysfunction action on hippocampal GABAergic interneurons (Schmid, et al., 2016; Haam & Yakel, 2017). Here, we have shown that a decrease in the conductance of cholinergic a7 nAChRs on OLM interneurons caused the impairment of induction of hippocampal synaptic plasticity.

## Acknowledgments

This research was supported by the Intramural Research Program of the NIH, National Institute of Environmental Health Sciences. IG and BSG acknowledge support from the Fondation pour la Recherche sur Alzheimer, CNRS, INSERM, ANR-17-EURE-0017 and ANR-10-IDEX-0001-02. BSG acknowledges funding from HSE Basic Research Program and the Russian Academic Excellence Project “5-100.”

## Supplementary Material

**Figure S1:**
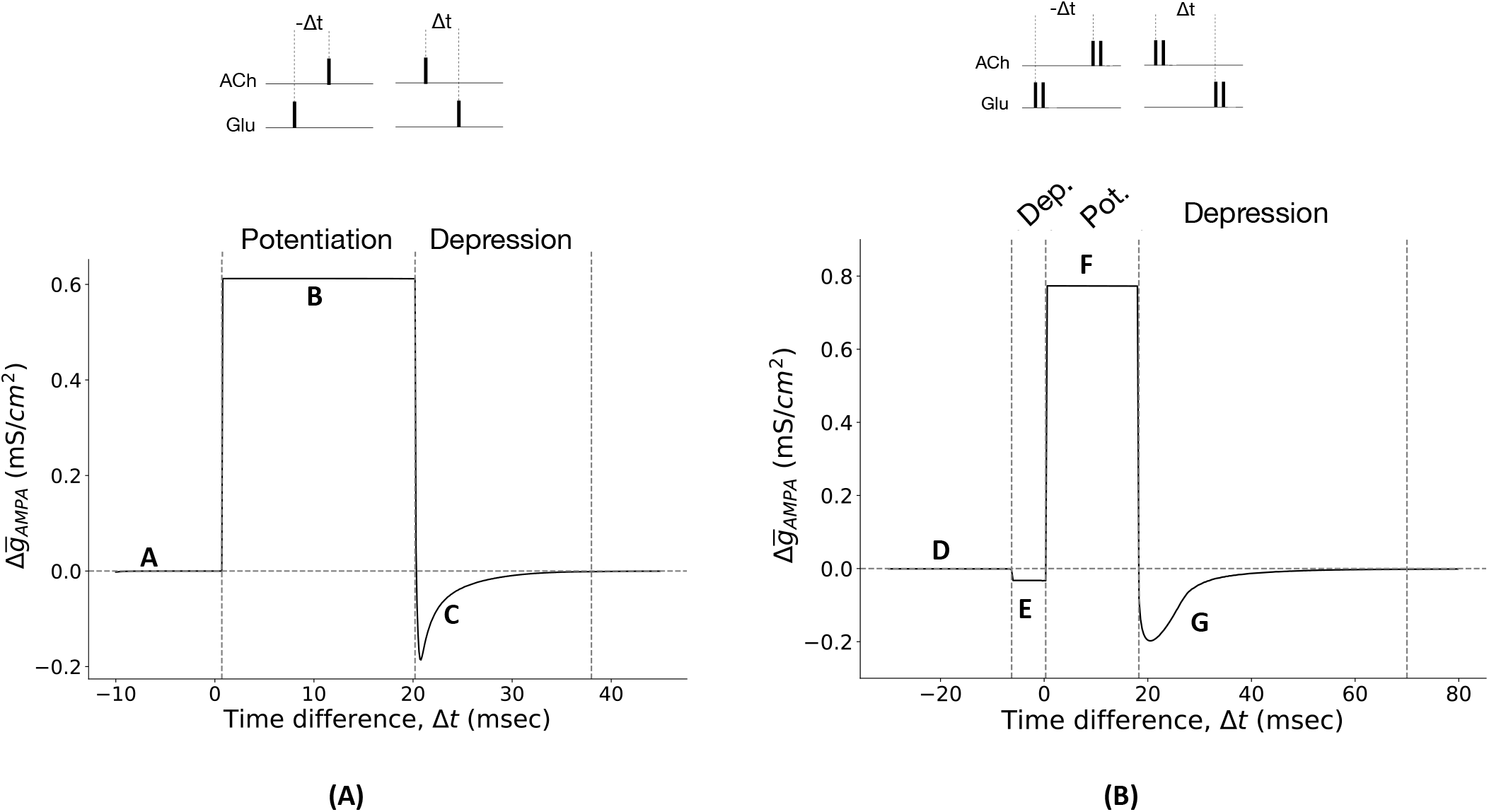
**(A)** Single spike pairs yield three qualitatively regions with respect to induction of plasticity. <u>Region A</u> (Δt < 0.7 msec): A pulse of glutamate activates the I-cell and E_D_. The I-cell releases one pulse of GABA that inhibits E_D_. In the meantime, the O-cell receives a pulse of ACh. Because the O-cell has a slower dynamic than the I-cell, by the time the O-cell releases GABA and inhibits the I-cell, the I-cells have already inhibited E_D_ and no plasticity is induced. <u>Region B</u> (0.7 < Δt < 20.2 msec): ACh activates the O-cell which inhibits the I-cell. The I-cell and E_D_ receive a pulse of glutamate with a delay Δt. By the time the I-cell and E_D_ receive the pulse of glutamate, the I-cell is in a hyperpolarized state and cannot release GABA. The feedforward inhibition is cancelled, i.e. no GABA arrives to E_D_, and the pulse of glutamate depolarizes E_D_, activating NMDARs and allowing enough calcium to enter the cell to induce potentiation. <u>Region C</u> (20.2 < Δt < 38 msec): As the arrival of the Glutamate pulse is delayed, eventually the I-cell will be able to spike and release one pulse of GABA. But because it is still recovering from the inhibition that received from the O-cell, GABA will arrive to E_D_ later than in Region A. E_D_ depolarizes just enough for calcium to cross the depression threshold before being inhibited, and depression is induced. **(B)** Single spike-douplets yield four qualitatively regions with respect to induction of plasticity. <u>Region D</u> (Δt < −6.28 msec): Similarly to Region A, glutamate is administrated before ACh. Two pulses of glutamate activate the I-cell, which releases 2 pulses of GABA that inhibit E_D_. ED cannot depolarize enough to activate enough NMDARs to induce plasticity. <u>Region E</u> (−6.28 < Δt < 0.3 msec): the I-cell receives 2 pulses of glutamate but only releases 1 pulse of GABA because of the inhibition is receiving from the O-cell. E_D_ depolarizes enough to allow calcium to cross the depression threshold, and depression is induced. Region F (0.3 < Δ*t* < 18.27 msec): I-cell receives 2 pulses of glutamate but it does not release any GABA because of the action of the O-cell. E_D_ depolarizes enough to allow calcium to cross the potentiation threshold and potentiation is induced. <u>Region G</u> (18.27 < Δ*t* < 70 msec): the I-cell releases either 1 or 2 pulses of GABA, depending on the delay between the 2 pulses of glutamate and the GABA released from the O-cell, which induces depression. The amplitude of depression depends on the either E_D_ received 1 or 2 pulses of GABA and the time at which these pulses arrive to E_D_.

**Figure S2:**
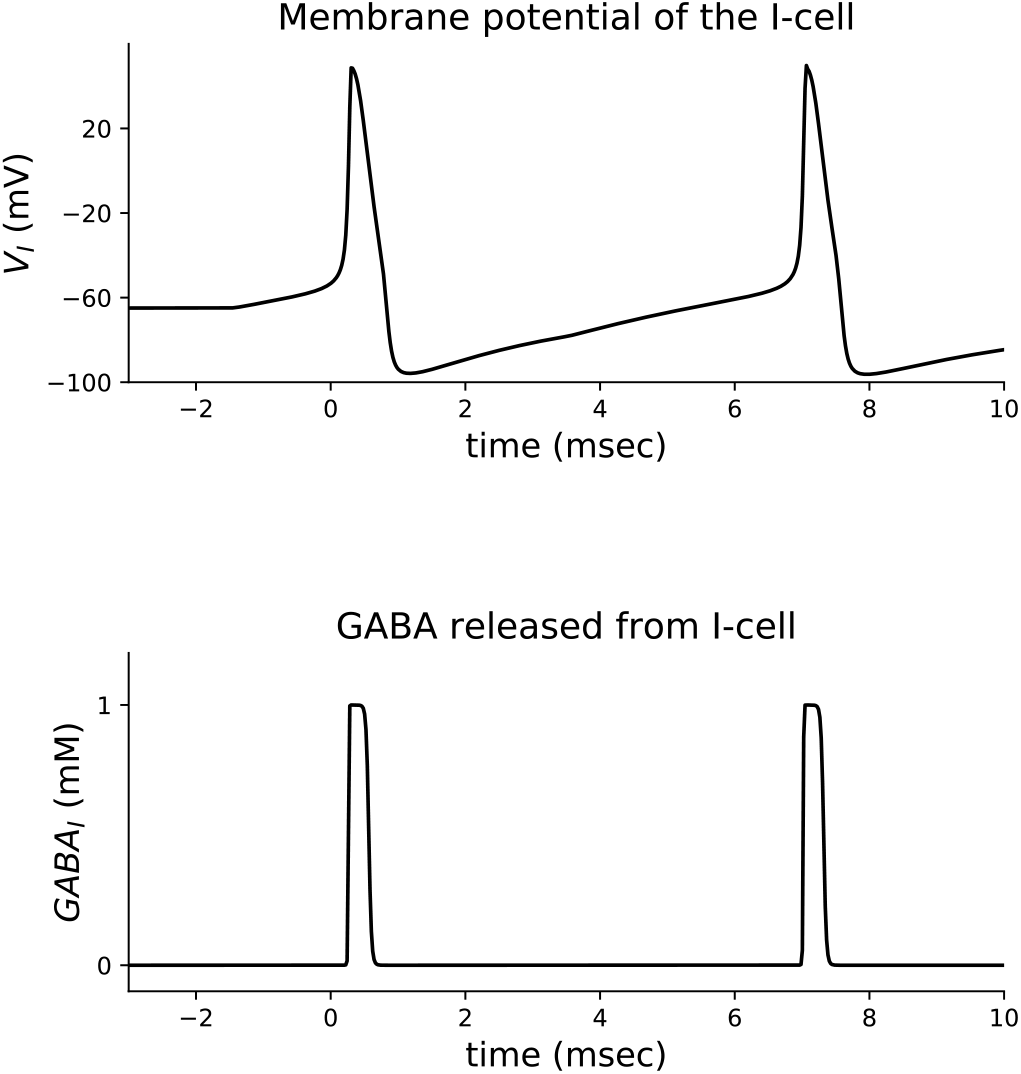
Membrane potential of the I-cell when it receives two pulses of glutamate (with amplitude 1mM and duration of 3 msec) with a frequency of 0.2 msec^-1^, and respective GABA release. GABA concentration can be described by a square function.

**Figure S3:**
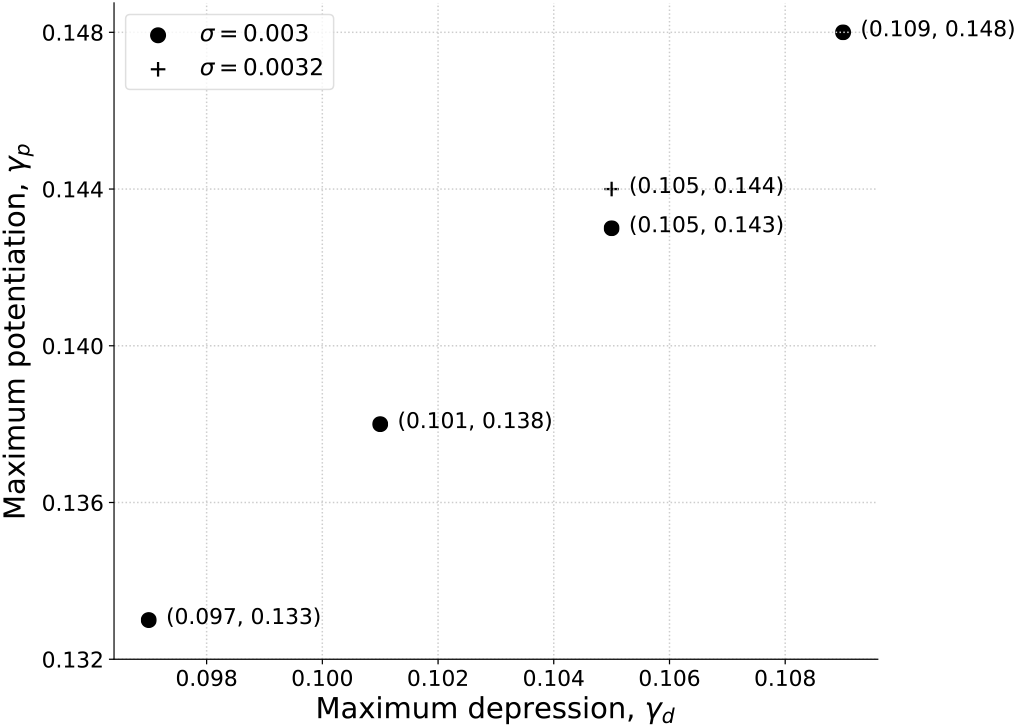
Set of parameters for which we obtain the same results as the ones shown in Fig.4.

**Figure S4:**
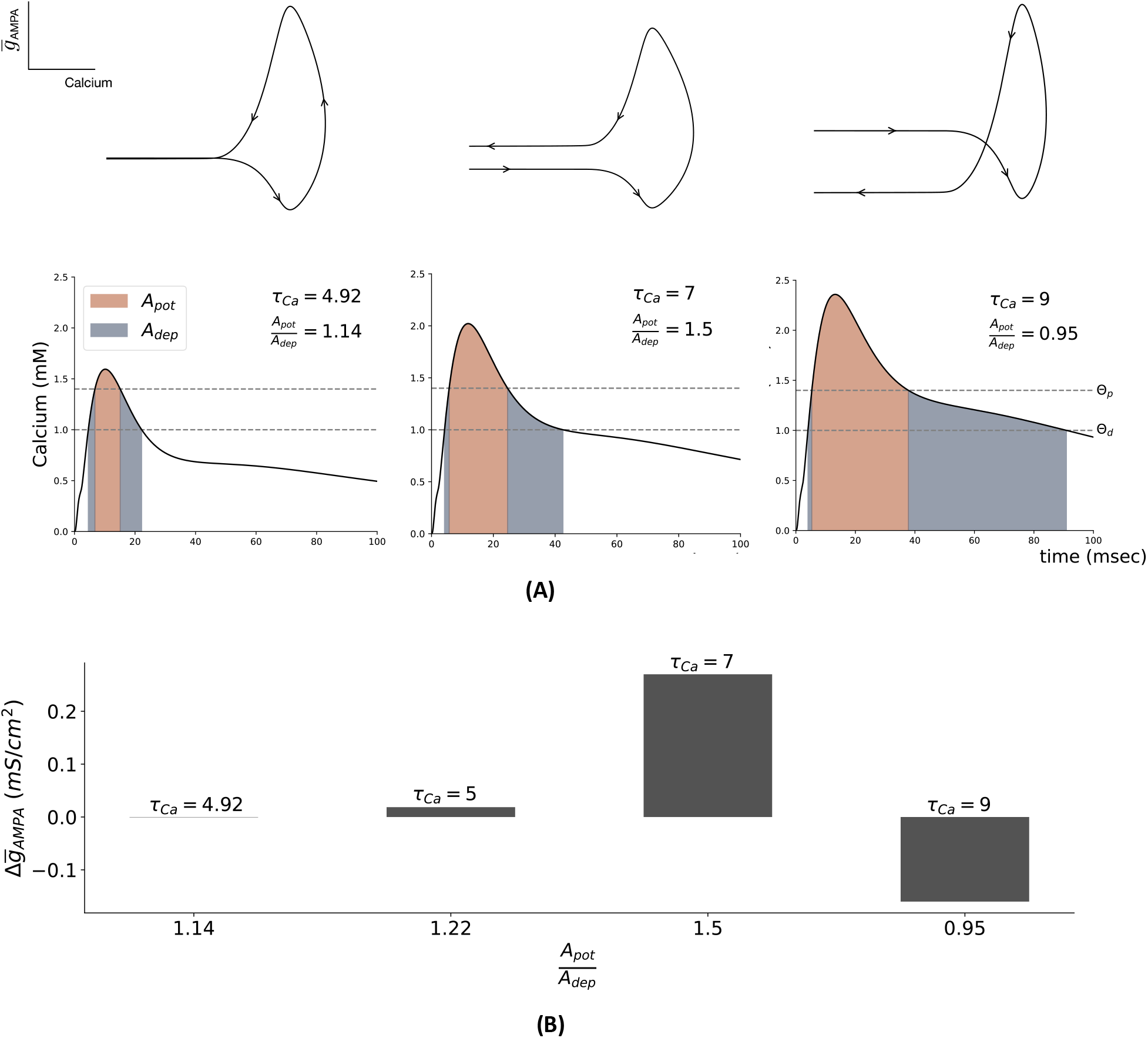
Increasing the ration between the area of potentiation and depression increases induction of synaptic plasticity. **(A)** By changing the calcium decay rate *τ*_Ca_, we change the ratio between the potentiation area (integral of calcium concentration above the potentiation threshold *θ*_p_) and the depression area (integral of calcium concentration above the depression threshold *θ*_d_ and bellow the potentiation threshold). **(B)** As the ratio between the potentiation and depression area increases, so does the induction of synaptic plasticity. In these numerical simulations, ED receives a pulse of glutamate followed by a pulse of GABA 2 msec after, each with an amplitude of 1mM and duration of 1 msec. The initial value of 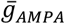 is 1.8 mS/cm^2^. All the other parameters are as indicated in Methods. The ratio between the potentiation and depression area is calculated by numerically integrating the area in the potentiation and depression zone, using the trapezoidal method.

